# Evaluation of growth and enzymatic characteristics of wild-type *Yarrowia lipolytica* strains

**DOI:** 10.64898/2026.03.28.715033

**Authors:** Idriss Ait-Tahar, Clémence Moret, Cécile Grondin, Alain Doyen, Eric Dugat-Bony, Catherine Madzak

## Abstract

*Yarrowia lipolytica* is a yeast of industrial interest exhibiting remarkable lipolytic and proteolytic capacities, with a high potential for white biotechnology applications. This yeast can be isolated from a wide range of natural, polluted or anthropogenic environments, including various food products. The present study aims to increase the data on *Y. lipolytica* phenotypic diversity by evaluating the growth parameters and secreted enzymatic activities of 28 wild-type *Y lipolytica* (and *Yarrowia* sp.) strains isolated from various environments across 10 countries. These data could facilitate the selection of appropriate strains for specific research purposes, particularly when wild-type strains are prioritized over genetically engineered ones, like for food-related applications. Notably, strain SWJ-1b exhibited an outstanding combination of favourable characteristics, with optimum (or near) performances for both growth and enzymatic parameters.

## Introduction

*Yarrowia lipolytica* is an oleaginous ascomycete yeast (class: *Saccharomycetes*, order: *Saccharomycetales*) that was noticed at first by some industrialists in the 1950s for its remarkable lipolytic and proteolytic capacities. According to the high levels of secreted lipase and protease activities of this yeast, wild-type *Y. lipolytica* strains were generally isolated from lipid-rich and/or protein-rich environments, notably from meat and dairy products (especially fermented ones, such as dry sausages and cheeses) and from sewage or oil-polluted waters and soils. Since a few decades, *Y. lipolytica* isolates were also found in a wider range of habitats, from marine waters, salt marshes, soils and rhizospheres to a variety of other food products (like fruits, vegetables or seafood) and even the faecal material of insects or vertebrates that consume them, as reviewed by Madzak (2021). This yeast species thus appears to exhibit a rather ubiquitous distribution, in both natural and man-made environments.

Besides the use of natural isolates, notably for single cell oil and protein production, as well as for depollution, *Y. lipolytica* is recognized since decades as a powerful host for heterologous protein expression, secretion and surface display (Madzak, 2021). The development of sequencing and genetic engineering tools, combined with an increasing knowledge of its metabolism, can now enable complex engineering of its metabolic pathways for numerous applications, and numerous laboratories throughout the world have chosen this yeast as chassis for microbial cell factories. White biotechnology applications of *Y. lipolytica* include notably single cell oil production, whole cell bioconversion and upgrading of industrial wastes, as reviewed recently (Madzak, 2021; Cao *et al*., 2023).

All research and applications involving genetically engineered *Y. lipolytica* strains need to comply with Genetically Modified Organisms (GMO) regulations, which vary between countries, especially concerning the release of GMOs. There are important differences notably between the USA and European countries. More specifically, the American regulatory approach places limited emphasis on production processes and instead relies on the concept of substantial equivalence whereas the European framework is more severe and holds to the precautionary principle (Madzak, 2021; Ricroch *et al*., 2026). If the problem of societal acceptance of GMO is particularly important for agriculture and agro-food industries, it also impacts negatively the domain of microbial biotechnology, while the corresponding legislations are still under construction in many countries (Ricroch *et al*., 2026). Consequently, some agro-industrial or commercial food companies are reconsidering their research and development strategies to favour the use of traditionally improved strains over that of GM ones in their production processes. Such societally driven strategical shifts have prompted a new interest in the study of the natural biodiversity of some yeasts, notably *Y. lipolytica* (Madzak, 2021). In this context, selecting appropriate *Y. lipolytica* strains from the numerous isolates available in international collections is a crucial step, whether for conventional or GMO-based research projects.

Among the many assets of *Y. lipolytica* for potential applications are its remarkable extracellular lipase and protease activities. Its sole extracellular lipase LIP2 is encoded by the eponymous gene, a member of the large *LIP* gene family (Fickers *et al*., 2011). *Y. lipolytica* secretes two proteases, the acid extracellular protease (AXP), encoded by the eponymous gene, and the alkaline extracellular protease (AEP), encoded by the *XPR2* gene. The expression and activity of these two proteases are tightly regulated by the pH of the growth medium. Their activities are largely mutually exclusive, with high AXP and low AEP activity at pH values below 6, when the opposite pattern is observed for pH values above 6 (Glover *et al*., 1997). These high levels of extracellular enzymes enable the growth of *Y. lipolytica* on a broad spectrum of substrates. Notably, protein- and/or lipid-rich agricultural and industrial by-products could be valorised as alternative substrates by selected *Y. lipolytica* strains, highlighting the strong potential of this yeast in white biotechnology applications (Madzak, 2021).

Although *Y. lipolytica* has a long history of use and has become a model yeast for studying lipid metabolism, its diversity remained poorly characterized until recently, as most studies focused on only a few laboratory strains. To address this limitation, Bigey *et al*. (2023) recently studied the genomic and phenotypic diversity of 56 wild-type *Y. lipolytica* strains isolated from diverse environments across 22 countries. Although *Y. lipolytica* strains are classified into five clades, no correlation was observed between clade distribution and the geographic or ecological origin of the isolates. Since *Y. lipolytica* exhibits a low genetic diversity (π = 0.0017) and possesses a pan-genome of 6,528 genes that differs only slightly from its core genome of 6,315 genes, this yeast appears to be a relatively recently evolving species, as reviewed by Bigey *et al*. (2023). In their study, 34 phenotypic traits were quantified and compared; however, establishing clear associations with specific genomic characteristics proved challenging. Nevertheless, 4 distinct mutational events in *XPR2* gene were shown to inactivate the corresponding AEP activity. Regarding lipid metabolism, most mutations were identified in genes belonging to previously described large gene families such as *ALK* or *LIP*, which likely limited their phenotypic impact. Consequently, the high diversity of lipid synthesis and accumulation in *Y. lipolytica* appears to be multifactorial and/or governed by complex regulatory mechanisms (Bigey *et al*., 2023).

As a model organism, particularly for the study of lipid metabolism, *Y. lipolytica* has had several of its strains fully sequenced and annotated, as recently reviewed (Madzak, 2021; Bigey *et al*., 2023). Almost all natural isolates of *Y. lipolytica* are haploid, carrying either a MATA or MATB allele, with the notable exception of the type strain CBS 6124 (referred to as CLIB 183 in this study) which is one of the rare diploid isolates. The first *Y. lipolytica* strain to be sequenced, CLIB 122 (also known as E150), was derived from a cross between the French strain W29 (CLIB 89) and the American strain CBS 6124-2 (CLIB 78) itself obtained by sporulation of the diploid type strain, followed by backcrosses and genetic engineering steps, as reviewed by Madzak (2021). CLIB 122 (E150) genome sequence and gene annotation still serve as the reference for this yeast species (Dujon 2004). By the early 2020s, nearly 30 wild-type or engineered *Y. lipolytica* strains had been sequenced (Madzak, 2021) and Bigey *et al*. (2023) subsequently performed the sequencing or resequencing of all 56 strains included in their study. As a result, genomic data are now available for 13 of the strains used in our work. These include some historically and industrially important strains such as W29 (CLIB 89), A101 (CLIB 82) and H222 (CLIB 80) as well as strains more recently sequenced by Bigey *et al*. (2023), namely CLIB 79, CLIB 200, CLIB 201, CLIB 202, CLIB 205, CLIB 206, CLIB 703, CLIB 791 (referred to as TL301 in this study), CLIB 879 (referred to as 141 in this study) and 1E07.

Building on the work published by Bigey *et al*. (2023), the present study aims to further expand the available data on *Y lipolytica* phenotypic diversity by evaluating the growth parameters and secreted enzymatic activities of 28 wild-type strains (including 2 *Yarrowia* sp.) isolated from various natural or polluted environments across 10 countries worldwide. These data are expected to provide the scientific community with valuable insights for selecting strains suited to specific research objectives, particularly when wild-type strains are favoured over genetically engineered ones, such as for food-related applications.

## Materials and Methods

### Strains

Twenty-eight *Y. lipolytica* (or *Yarrowia* sp.) wild strains (**Table 1**) were selected either among the numerous wild-type strains kept at CIRM-Levures (https://cirm-levures.bio-aware.com) INRAE’s yeast collection (23 strains), or from the dairy strains collection of the SayFood laboratory (5 strains). All strains were identified as *Y. lipolytica* at CIRM-Levures or SayFood laboratory (unpublished data) with the exception of ME-37 (CLIB 3560) and ME-54 (CLIB 3561) which were assigned only to the genus *Yarrowia*.

**Table 1.** Characteristics of *Yarrowia lipolytica* (or *Y. species*) strains used in this work.

| Strain name<br>(this study) | CIRM *<br>reference | Other name(s) | Ploidy<br>(Mat locus) | Source<br>category | Description<br>(Town or Region,<br>Country) | Bioproject<br>accession<br>number | *Y.*<br>*lipolytica*<br>clade | Main<br>references |
| --- | --- | --- | --- | --- | --- | --- | --- | --- |
| H222 | CLIB 80 | NA | haploid<br>(Mat A) | soil | isolated from soil<br>(Germany) | PRJEB28424 | Clade 1 | Barth & Weber, 1983<br>Madzak, 2021 |
| CLIB 202 | CLIB 202 | NA | haploid<br>(Mat A) | soil | isolated from soil<br>(Netherlands) | PRJEB42834 | Clade 4 | Naumova *et al.*, 1993 |
| CLIB 205 | CLIB 205 | NA | haploid<br>(Mat A) | soil | isolated from soil<br>(Russia) | PRJEB42834 | Clade 2 | Naumova *et al.*, 1993 |
| CLIB 206 | CLIB 206 | ATCC 20226,<br>CBS 7033 | haploid<br>(Mat B) | soil | isolated from soil<br>(Japan) | PRJEB42834 | Clade 4 | Naumova *et al.*, 1993 |
| CLIB 703 | CLIB 703 | ATCC 48436,<br>CBS 6303,<br>JCM 8054,<br>NBRC 10073 | haploid<br>(Mat A) | soil | isolated from soil<br>(Japan) | PRJEB42834 | Clade 3 | Yamada & Machida,<br>1963<br>Madzak, 2021 |
| W29 | CLIB 89 | ATCC 20460,<br>CBS 7504,<br>CICC 1778,<br>NBRC 113670,<br>NRLL Y-3178,<br>VKPM Y-3178 | haploid<br>(Mat A) | polluted<br>environment | isolated from<br>a sewer<br>(Paris, France) | PRJNA295780 | Clade 3 | Gaillardin *et al.*, 1973<br>Madzak, 2021 |
| A-101 | CLIB 82 | NA | haploid<br>(Mat B) | polluted<br>environment | isolated in a<br>car wash (Poland) | PRJEB14097 | Clade 2 | Wojtatowicz *et al.*,<br>1991<br>Madzak, 2021 |
| SWJ-1b | CLIB 3761 | MCCC 2E00068 | haploid | animal | isolated from the gut<br>of a marine fish<br>(Bohai sea, China) | NA | NA | Chi *et al.*, 2007<br>Madzak, 2021 |
| ME-37<br>(*Y. species*) | CLIB 3560 | NA | haploid | animal | isolated on a *Hoplia*<br>*coerulea* beetle<br>(Occitanie, France) | NA | NA | NA |
| ME-54<br>(*Y. species*) | CLIB 3561 | NA | haploid | animal | isolated on a *Hoplia*<br>*coerulea* beetle<br>(Occitanie, France) | NA | NA | NA |
| CLIB 183 | CLIB 183 | ATCC 18942,<br>YB-423,<br>CBS 6124,<br>JCM 2320,<br>JCM 8060,<br>MUCL 29853,<br>NBRC 1548,<br>NRRL YB-423 | diploid<br>(MatA,<br>MatB)<br>[type strain] | agro-food | isolated in a<br>maize-processing<br>plant (USA) | NA | NA | von Arx, 1972<br>Madzak, 2021 |
| CLIB 79 | CLIB 79 | CBS 6125 | haploid<br>(Mat A) | agro-food | isolated in a<br>maize-processing<br>plant (USA) | PRJEB42834 | Clade 3 | Neuvéglise *et al.*,<br>2005 |
| EL13-B1-2-1 | CLIB 3040 | NA | haploid | food | isolated from<br>refrigerated<br>vegetables (France) | NA | NA | NA |
| CLIB 200 | CLIB 200 | CBS 599,<br>DBVPG 6132,<br>IFO 1195,<br>VKM Y-47,<br>BCRC 20864 | haploid<br>(Mat A) | food | isolated from<br>rancid margarine<br>(Netherlands) | PRJEB42834 | Clade 4 | Naumova *et al.*, 1993 |
| CLIB 201 | CLIB 201 | CBS 2074,<br>DBVPG 6022 | haploid<br>(Mat A) | food | isolated from olives<br>(Italie) | PRJEB42834 | Clade 1 | Naumova *et al.*, 1993 |
| ME-674 | CLIB 3752<br># | NA | haploid | food | isolated from<br>breadmaking leaven<br>(Grand-Est, France) | NA | NA | NA |
| CAM-25-E | NA | NA | haploid | dairy/cheese | isolated from a dairy<br>product processing<br>line (France) | NA | NA | NA |
| CAM-28-E | NA | NA | haploid | dairy/cheese | isolated from a dairy<br>product processing<br>line (France) | NA | NA | NA |
| CAM-43-C | NA | NA | haploid | dairy/cheese | isolated from a<br>Camembert cheese<br>(France) | NA | NA | NA |
| 1E07 | NA | NA | haploid<br>(Mat B) | dairy/cheese | isolated from a<br>Livarot cheese<br>(France) | PRJEB42834 | Clade 4 | Mansour *et al.*, 2009 |
| Ex-fcom-LP | NA | NA | haploid | dairy/cheese | isolated from an<br>Epoisses cheese<br>(France) | NA | NA | NA |
| TL301 | CLIB 791 | NA | haploid<br>(Mat A) | dairy/cheese | isolated from a<br>"chevrotin des Aravis"<br>goat cheese<br>(Haute-Savoie, France) | PRJEB42834 | Clade 1 | Adouard *et al.*, 2015 |
| TL302 | CLIB 632 | NA | haploid | dairy/cheese | isolated from a<br>"chevrotin des Aravis"<br>goat cheese<br>(Haute-Savoie, France) | NA | NA | Adouard *et al.*, 2015 |
| R18-455 | CLIB 3749<br># | NA | haploid | dairy/cheese | isolated from a goat<br>cheese (Takamaka,<br>Réunion, France) | NA | NA | NA |
| R18-456 | CLIB 3750<br># | NA | haploid | dairy/cheese | isolated from a goat<br>cheese (Takamaka,<br>Réunion, France) | NA | NA | NA |
| R18-457 | CLIB 3751<br># | NA | haploid | dairy/cheese | isolated from a goat<br>cheese (Takamaka,<br>Réunion, France) | NA | NA | NA |
| 141 | CLIB 879 | NA | haploid<br>(Mat A) | dairy/cheese | isolated from a<br>pressed sheep cheese<br>(Navarre, Spain) | PRJEB42834 | Clade 1 | NA |
| 296 | CLIB 880 | NA | haploid | dairy/cheese | isolated from a<br>pressed sheep cheese<br>(Navarre, Spain) | NA | NA | NA |

\*: CIRM-Levures online catalog link: https://cirm-levures.bio-aware.com/. #: recent CIRM-Levures acquisition; addition to the website in progress. NA: not available.

### Growth conditions and determination of growth parameters

All strains were maintained on Yeast-Peptone-Dextrose (YPD) plates (1% yeast extract, 1% bacto peptone, 1% glucose, 1% Agar) incubated at 28°C and pre-cultivations were performed in 2 mL YPD liquid medium incubated overnight at 28°C in shake-flasks at 180 rpm. Growth parameters in rich YPD medium were determined using an Agilent BioTek H1 automated microplate reader, following manufacturer’s specifications, and Greiner 96 Black Flat Bottom Fluotrac microplates with lids. For each experiment, all wells from a 96-wells microplate received 200 μL, of YPD inoculated with *Y. lipolytica* cells at an initial OD_600_ of 0.01 for the 60 inner wells, or of YPD as blank control for the external wells. Microplates were incubated at 28°C with continuous shaking at 731 rpm for 24-48h (until growth reached a plateau) and OD_600_ readings were recorded hourly. Growth parameters, presented in **Figure 1**, were calculated from the acquired data using BioTek Gen5 software (version 2.06) as illustrated in **Figure S1**. Each experiment included at least 3 independent biological replicates, each with a minimum of 6 technical replicates, for each strain.

**Figure 1.**
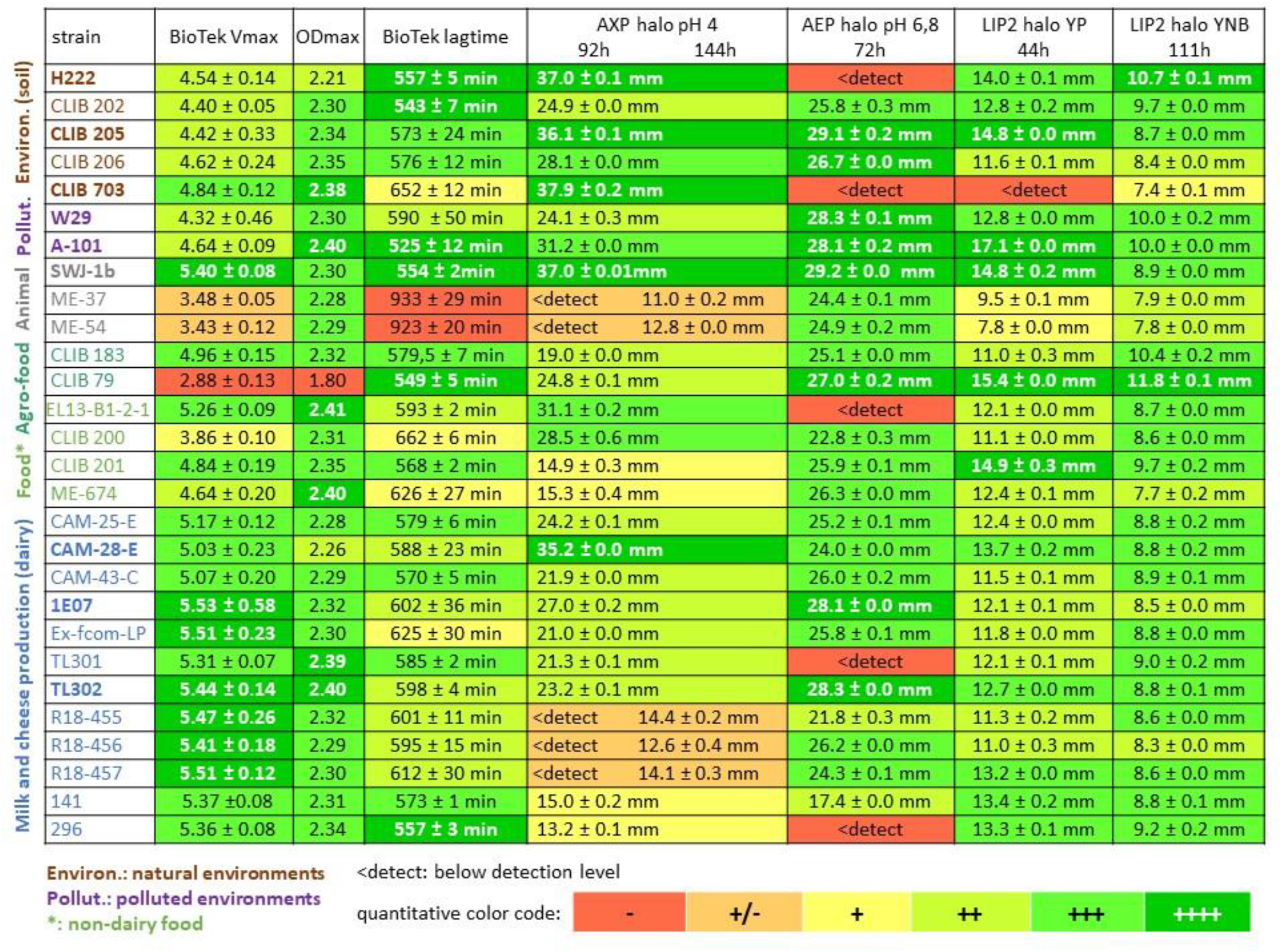
Phenotypic characteristics of the 28 *Y. lipolytica* or *Y*. sp. strains used in this work. This Figure presents, in the form of a color-coded Table, all the data recovered from the strains of the present study: growth parameters extracted from BioTek analysis (Vmax, ODmax and lagtime, as illustrated in Figure S1), AXP and AEP protease activities (halo diameter on skim milk plates) and LIP2 lipase activities on rich and minimal medium (halo diameter on tributyrin plates, as illustrated in Figure S2). A quantitative colour code was used to improve data readability ranging from the least (-) favourable results (highest lagtime; lowest Vmax, ODmax or activities) in red, to the most (++++) favourable results (lowest lagtime; highest Vmax, ODmax or activities) in dark green. The 28 strains were grouped by ecosystems categories indicated by color coding: natural environments (soils) in brown; polluted environments in purple; animals in grey; non-dairy food and agro-food in green (light and dark, respectively) and dairy environments (milk and cheese production) in blue. The nine strains exhibiting the most notable combinations of characteristics, or the highest performance for at least one enzymatic activity, have their names highlighted in bold.

### Measurement of extracellular LIP2 lipase activity

LIP2 activity was assessed by measuring the diameter of halos formed on either rich (YP: 1% yeast extract, 1% bacto peptone, 1% Agar) or minimal (YNB: 0.17% Yeast Nitrogen Base, 1% Agar) medium plates containing 1 % tributyrin as main carbon source, as previously described (Bigey *et al*., 2023). In both cases, plates were inoculated with patches of 10^5^ cells (10 μL at OD_600_ 0.3) and incubated at 28 °C, with halo development assessed daily. Halo diameters were measured after 44h for YP-based plates and 111h for YNB-based plates. Measurements were performed in triplicate (technical repeats) on three halos per strain (biological repeats), using ImageJ software (Schneider *et al*., 2012) on plate photographs, with the plate diameter being used for calibration, as illustrated in **Figure S2**.

### Measurement of extracellular AEP and AXP protease activities

As AEP and AXP, the two *Y. lipolytica* secreted proteases, are mainly expressed and active respectively above and below pH 6 (Glover *et al*., 1997), their activities were assessed by measuring the diameter of halos formed on skim milk plates, as previously described (Bigey et al., 2023), but with the addition of buffering solutions for pH control. For AEP, plates contained 2% skim milk, 0.17% yeast nitrogen base, 0.1% glucose, 1% agar, and 50 mM phosphate buffer at pH 6.8. For AXP, plates contained the same components but with 50 mM citrate buffer at pH 4. In both assays, plates were inoculated with patches of 10^5^ cells and incubated at 28 °C, with halo development assessed daily. Halo diameters were measured after 72h for AEP and after 92h or 144h for AXP in triplicate (technical repeats) on three halos per strain (biological repeats), using ImageJ software (Schneider *et al*., 2012) on plate photographs, with the plate diameter used for calibration, as described for the lipase assays.

## Results

The wild-type *Y. lipolytica* (or *Yarrowia* sp.) strains selected for this study have been chosen from the CIRM-Levures and SayFood collections as samples as wide as possible of *Y. lipolytica* biodiversity in terms of environments and countries, as shown in **Table 1**. These 28 strains can be grouped into 6 categories *i*.*e*. natural environments (soils, 5 strains), polluted environments (2 strains), animals (3 strains), agro-food industry (2 strains), non-dairy food (4 strains) and dairy products (milk or cheese production, 12 strains). The ME-37 and ME-54 strains, although identified only as *Yarrowia* sp., were nonetheless included in our study due to their unusual source, having been isolated from the cuticle of an insect species. For each strain, the parameters of growth during cultivation in microplates on YPD medium (lagtime, Vmax and ODmax), as well as the main secreted enzymatic activities (LIP2 lipase, AXP and AEP proteases) were determined, as shown in **Figure 1**. The use of a color code, from red (for less favourable results) to dark green (for more favourable ones) allows to visualize more clearly the main results.

Each of the characteristics tested presents some diversity between strains, with the AEP activity (proteolytic activity at pH value above 6) being the more variable and the LIP2 activity in minimal (YNB) medium being the less variable. The variability of these characteristics cannot be fully linked to particular ecosystems, since there is a large range of performance levels for most parameters among the strains isolated from similar ecosystems, notably visible for the soil, “non-dairy food / agro-food” or dairy categories that contain the more strains.

Nevertheless, a few global tendencies can be observed: environmental strains (soils, polluted environments) appear to present a lower Vmax but shorter lagtime, when the contrary is seen for dairy strains; they also seem to exhibit higher secreted proteolytic and lipolytic activities, despite a few exceptions.

Nine strains, distributed across the main different ecosystems, exhibiting either a remarkable combination of traits or an optimal performance for at least one enzymatic activity, are highlighted in bold in **Figure 1**. Among them, the animal-derived strain SWJ-1b stands out as a “super-strain”, displaying optimal or near-optimal performances across all evaluated parameters. Only three additional strains combined robust growth with both high proteolytic and lipolytic activities i.e. the soil-derived strain CLIB 205, the polluted-environment strain A-101 (CLIB 82) and the dairy strain CAM-28-E.

Some other strains exhibited robust growth and remarkable performances, but only for some of the evaluated activities. In particular, the highest Vmax values were observed for the two dairy strains 1E07 and TL302 (CLIB 632), which also exhibit a high AEP activity. The polluted-environment strain W29 (CLIB 89) similarly showed a high AEP activity along with a high LIP2 activity. The soil-derived strain H222 (CLIB 80) demonstrated both high AXP and high LIP2 activities although its AEP activity is undetectable. Finally, the soil-derived strain CLIB 703, exhibited the highest AXP activity, but lacked both AEP and LIP2 activities.

The only diploid strain, CLIB 183, did not particularly stand out, either in terms of growth parameters or secreted enzymatic activities, displaying only moderate characteristics. The two *Yarrowia* sp. strains (ME-37 and ME-54) exhibited a very unusual growth pattern, characterized by an exceptionally long lag phase combined with a low Vmax.

Well-isolated clonal colonies of each of these nine strains were grown on YPD plates for 4 days and photographed under standardized conditions to allow comparison of colony sizes and filamentation (**Figure 2**). While minor differences in colony diameter and color were observed among the 9 strains, the most striking differences concerned the degree of filamentation. All environmental strains, from both soils and polluted environments, displayed mild to strong filamentation, whereas the fish-derived SWJ-1b and all the dairy strains formed smooth colony with almost no sign of dimorphic transition. Despite sharing a filamentous phenotype, the five environmental strains exhibited marked differences in colony morphology, illustrating how the timing and intensity of dimorphic transitions can influence the final colony shape.

**Figure 2.**
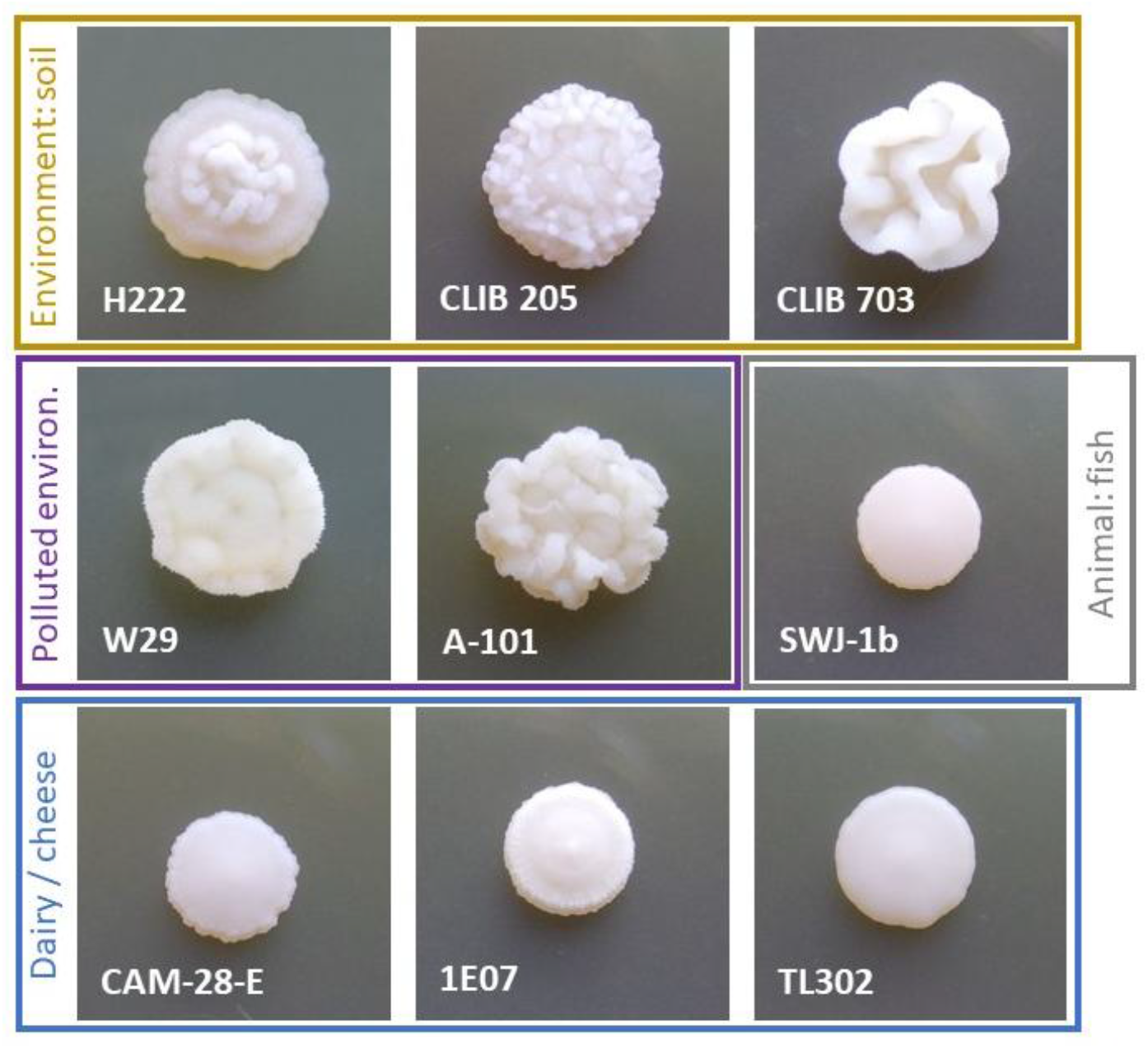
Visual aspect of selected *Y. lipolytica* clonal colonies on rich medium. The colony shown are representative of the typical appearance for each strain. Isolation source categories are indicated with the same colour code as in previous figure.

## Discussion

By analysing the growth parameters and secreted enzymatic activities of 28 wild-type *Y lipolytica* (and *Yarrowia* sp.) strains isolated from diverse environments across 10 countries, our study identified the fish-derived SWJ-1b as a “super-strain” exhibiting optimal, or near optimal, performances for all the tested characteristics. SWJ-1b was originally isolated from the gut of a marine fish from Bohai Sea (China) by the Ocean University of China (OUC, Qingdao, PR China; Chi *et al*. 2007). This strain is notable for its high protein content and has been applied to white biotechnology applications (Madzak, 2021) through genetic engineering (Cui *et al*. 2011; Tan *et al*. 2016) or through the development of non-GM mutant strains (Liu *et al*. 2017). In addition to SWJ-1b, eight other strains, including six also available from CIRM-Levures, display robust growth characteristics and interesting levels of secreted activities (**Figure 1**). Among them, only three strains cumulate both high proteolytic and lipolytic activities: CLIB 205 (highest AEP activity, with SWJ-1b), A-101 (CLIB 82) and CAM-28-E. The A-101 strain has notably been applied to *in situ* soil bioremediation and, through genetic engineering or through selection or generation of mutant strains, to white biotechnology applications (Madzak, 2021). The other five strains exhibited high performance only for specific secreted activities. More specifically, CLIB 703 exhibited the highest AXP activity (proteolytic activity at pH value below 6), H222 (CLIB 80) demonstrated high AXP and lipolytic activities, W29 (CLIB 89) showed high AEP (proteolytic activity at pHs above 6) and lipolytic activities and the two dairy strains 1E07 and TL302 (CLIB 632), were associated to a high AEP activity with the highest Vmax. Therefore, these data could be helpful in the selection of a strain, depending on the type of secreted activity and the pH conditions required. For project involving genetic engineering, auxotrophic derivatives (notably Ura^-^) of five of these nine remarkable strains (H222, W29, A-101, SWJ-1b and 1E07) are already available (Madzak, 2021). Notably, the W29 strain had been genetically engineered since several decades to generate the series of Po1a to Po1z GM strains, applicable to heterologous expression and protein production for various white biotechnology applications (Madzak, 2021; Moret & Madzak, 2025).

Despite some overall trends observed in the data from **Figure 1**, notably that most environmental strains exhibited higher secreted proteolytic and lipolytic activities, the nine most notable strains were distributed across different ecosystem types, with the exception of the “non-dairy food / agro-food” category. The six strains from this category generally lacked remarkable characteristics, except for CLIB 79, which displayed high AEP and lipolytic activities. However, this strain performed poorly in growth parameters, displaying the lowest Vmax and ODmax. This counter-performance could be explained by the fact that CLIB 79, isolated in a maize-processing plant, could be better adapted to growth at lower temperatures (refrigerated environments) whereas our cultivations were conducted at 28 °C. The nine most notable strains were also distributed across four different *Y. lipolytica* clades, indicating that they are not closely genetically related. The two insect-derived *Yarrowia* sp. strains ME-37 and ME-54 exhibited highly unusual growth patterns, confirming that they do not belong to *Y. lipolytica* species. Their precise taxonomic affiliation remains to be determined through genome analysis at CIRM-Levures.

Thirteen strains from our study, H222, W29, A-101, CLIB 79, CLIB 200, CLIB 201, CLIB 202, CLIB 205, CLIB 206, CLIB 703, 1E07, TL301 (CLIB 791) and 141 (CLIB 879), were also included in the biodiversity study on 56 strains performed by Bigey *et al*. (2023), allowing a comparison with our results on some parameters. While the two studies agree on some points, direct comparisons are challenging due to methodological differences. Notably, protease assays in the 2023 study were performed on non-buffered skim milk agar (SMA) for which the pH was not reported whereas our study used two types of buffered skim-milk-based plates to discriminate between AXP (proteolytic activity at pH 4) and AEP (proteolytic activity at pH 6.8). Despite this lack of pH discrimination, the absence of proteolytic activity in H222, CLIB 703 and TL301 (CLIB 791) strains in the study of Bigey et al. (2023) correlates with the lack of AEP activity observed in our study, suggesting that the pH of their SMA plates was likely above 6.

The 2023 study was able to link the lack of proteolytic activity in some strains to genetic modifications in *XPR2* gene (for example the Ser196Tyr point mutations in H222 and the stop codon 60 in TL301) whereas the deficiency in other strains, such as CLIB 703, remained unexplained. Regarding lipase activity, both studies used YP- or YNB-based plates but the measurement methodologies differed. Indeed, whereas our study measured halo diameters from standardized inoculations (**Figure S2**), Bigey *et al*. (2023) measured only the thickness of halos around colonies, yielding smaller and likely less significant values. This difference could explain some observed discrepancies, such as strains for which we detected satisfactory LIP2 level, notably CLIB 79, CLIB 200, CLIB 202, 1E07 and 141 (CLIB 879) on both media, and H222, CLIB 703 and TL301 (CLIB 791) on YNB-based tributyrin medium, despite the 2023 study reporting null or very weak lipase activity. For instance, the robust LIP2 activity of CLIB 200 on YNB-based plates is evident in Figure S2, whereas the previous study reported no detectable lipase activity for this strain (Bigey *et al*., 2023).

Besides these few discrepancies, our study confirmed some of the conclusions of Bigey *et al*. (2023), notably the high diversity of metabolic phenotypes and morphotypes among isolates and the difficulty precisely linking these phenotypic characteristics to environments or geographic origins. In conclusion, the data obtained are expected to inform the scientific community in the selection of suitable strains for targeted research applications. Notably, they could allow the identification of *Y. lipolytica* strains with robust growth and the high levels of secreted pH-adapted proteolytic and/or lipolytic activities enabling the valorization of protein- and/or lipid-rich agricultural and industrial byproducts as innovative substrates for white biotechnology applications.

## Acknowlegments

This research was conducted with the support of the Department of Cooperation and Cultural Affairs of the Consulat Général de France in Québec (Samuel de Champlain partnership, convention n° 2021/013). The authors thank Prof. Zhenming Chi (OUC, Qingdao, PR China) and the MILA culture collection (Laboratoire des Micro-organismes d’Intérêt Laitier et Alimentaire, Université de Caen, France) for kindly providing the SWJ-1b and 1E07 strains, respectively.

