## Supplementary figures and images for "Evaluation of growth and enzymatic characteristics of wild-type *Yarrowia lipolytica* strains"

### Suplemental Figure S1

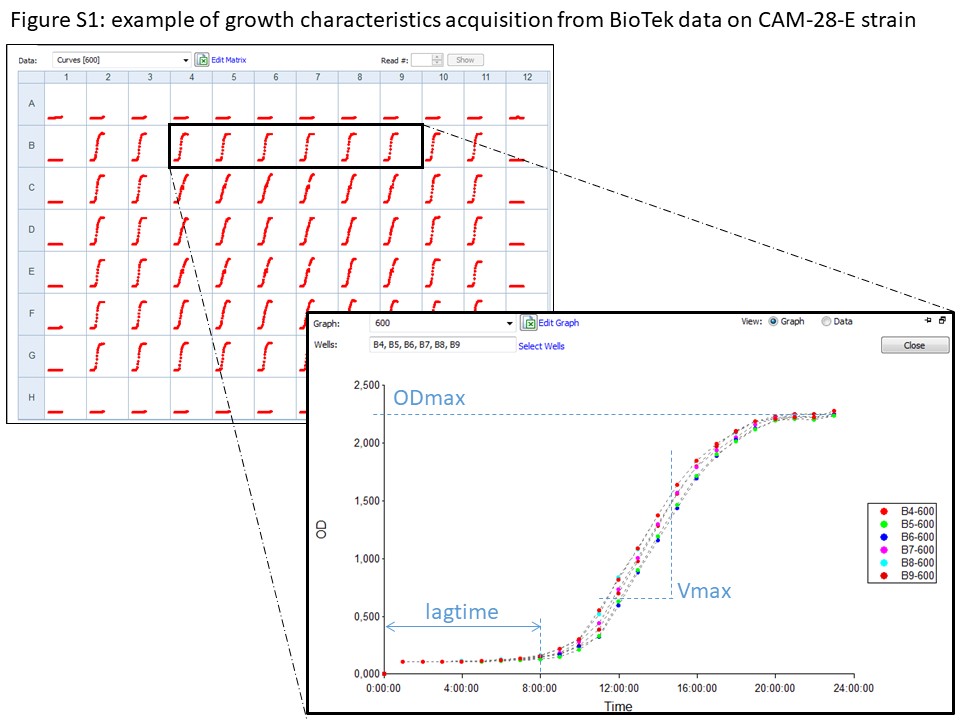

### Suplemental Figure S2

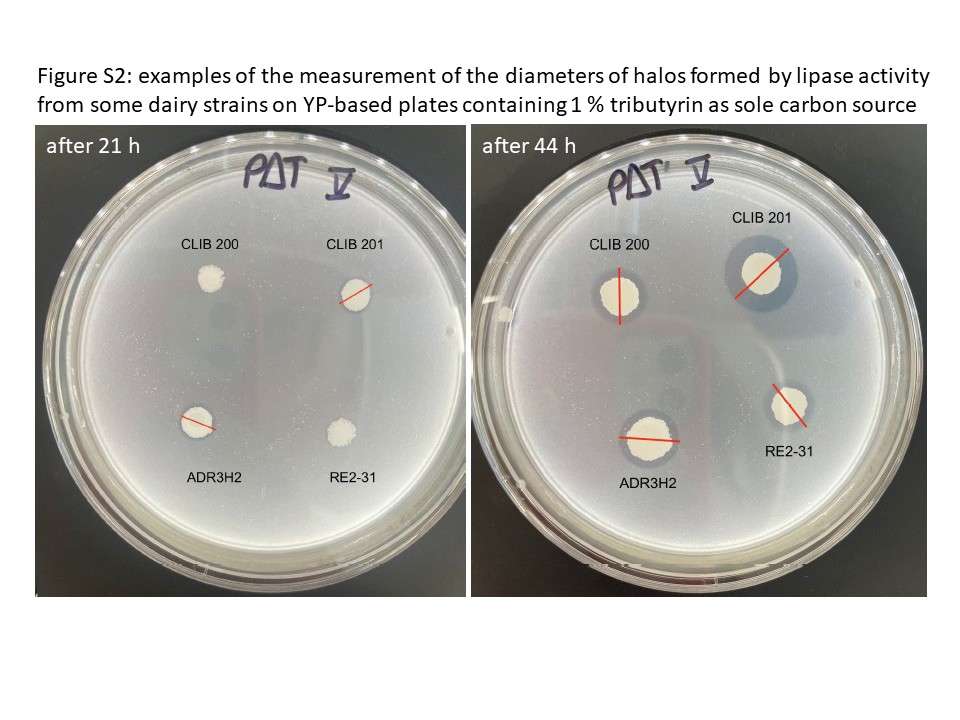
